# Pleiotropic antimalarial activities and immunomodulation exhibited by Himalayan Buransh (*Rhododendron arboreum*) in human and rodent malaria models

**DOI:** 10.64898/2026.06.01.729236

**Authors:** Cherish Prashar, Nishtha Tiwari, Reva S. Thakur, Sakshi Anand, Renuka Harit, Ruby Bansal, Ahmed Mohsin, Priyanka Rani, Hem Lata Singh, Padam Raj Bhatt, Himanshu Kumar, Vineeta Singh, Soumyananda Chakraborti, Rajesh Kumar Joshi, Brijesh Rathi, Jyoti Das, Mohammad Abid, Shailja Singh, Amol Gurav, Kapil Vashisht, Kailash C. Pandey

## Abstract

Emerging drug resistance against the malaria parasite is worrisome and necessitates the development of novel antimalarials. Himalayan Buransh (Rhododendron arboreum) is a well-known medicinal plant found in the northern states of India. In this study, we observed the pleiotropic antimalarial activities and immunomodulation exhibited by the aqueous extract of Buransh flower (AEBF). AEBF demonstrated significant IC50 values (16–29 µg/ml) against the asexual stages of various *P. falciparum* strains (3D7, Dd2-chloroquine-resistant and C580Y-artemisinin resistant). The oral administration of AEBF (200 mg/kg) in mice, suppressed ∼80% *P. berghei* parasitemia, improved mean survival time (MST-23.5 days) and prevented splenomegaly. Notably, the combination of AEBF and artesunate not only cleared primary infection, but also conferred sustained immunity. This immunomodulatory effect, driven by protective IFN-γ resulted in reduced parasitemia during a homologous challenge without the need for further treatment. It is important to highlight the malaria transmission blocking activity of AEBF, resulting in reduced sexual stage male gametocyte exflagellation. Furthermore, the virtual drug screening of selected bioactive constituents from Buransh flower demonstrated potent binding against multiple *P. falciparum* proteins, suggested a pleiotropic mode of action. Altogether, our results corroborated the first ever evidence of the multistage antimalarial potential of Buransh flower, supported by in vitro cell studies, in vivo rodent malaria model and in silico docking analyses. Based on our study’s findings and the traditional use of Buransh juice as a medicinal beverage in Uttarakhand, India, we propose exploring it as an adjunct therapy for drug-resistant malaria, subject to further clinical validation.

## Introduction

The global fight against malaria is under serious threat due to increasing drug resistance. The World Malaria Report 2023 highlighted a significant rise in treatment failures for *P. falciparum* malaria in the WHO African Region between 2015 and 2022, indicating the spread of resistance to Artemisinin Combination Therapies (ACTs) including the partner drugs . This alarming trend extends to other antimalarial drugs, as resistance is emerging across various malaria-endemic regions (Atul et al., 2023; Das et al., 2021; Mukherjee et al., 2017; Ndwiga et al., 2021; Nima et al., 2022; Takala-Harrison et al., 2015). Given the absence of an effective replacement for artemisinin and its derivatives, it has become crucial to explore alternative or supplementary treatments to combat the growing challenge of drug resistance.

Nature-derived chemical scaffolds had been the cornerstone for the development of antimalarials (Prashar et al., 2022). Rhododendron of the *Ericaceae* family is found in various regions, including Northern hemisphere, Southeast Asia, North America, Europe, and Northeastern Australia (Guo et al., 2017; Qiang et al., 2011). *Rhododendron arboreum*, commonly known as “Buransh” in the Uttarakhand region of northern India, is a plant with distinctive red flowers found in the western and eastern Himalayas. Due to its economic importance Buransh has been awarded a Geographical Identification: 866 by the Department of Industry Promotion and Internal Trade, Ministry of Commerce and Industry, Government of India. Further on in the manuscript, we will refer *Rhododendron arboreum* as “Buransh”. In the Uttarakhand region of India, the bright red flowers of this plant are used to make a popular local beverage called ‘Buransh juice’, widely enjoyed by local communities.

Numerous studies have explored the diverse therapeutic potential of Buransh. Various parts of the plant, including leaves, flowers, stem, bark, and roots, have exhibited a range of biological activities. The leaves are rich in bioactive compounds such as alkaloids, terpenoids, flavonoids, tannins (Dai et al., 2021), and have shown anti-inflammatory, antioxidant, antifungal, and cardiovascular protective properties. Buransh juice is often consumed to alleviate high-altitude sickness and digestive disorders (Dai et al., 2021); while the bark has demonstrated anti-cancer, anti-mutagenic, and anti-microbial effects (Hariharan and Rangaswami, 1966; Nisar et al., 2013). Given these impressive biological activities, it is important to investigate the potential of Buransh to combat pathogens such as *Plasmodium*, the primary cause of malaria in Southeast Asia and Sub-Saharan Africa. Notably, a recent Canadian study reported the anti-plasmodial properties in essential oils from *Rhododendron subarcticum* leaves, but these oils demonstrated cytotoxicity in human red blood cells (Séguin et al., 2023). This underscores the need for safer, plant-based anti-malarial alternatives. The present study explored the multistage antimalarial effects of Aqueous Extract of Buransh Flower (AEBF) using in vitro cell studies, in vivo therapeutic evaluation in mice and molecular docking analyses.

## Materials and methods

### Preparation of aqueous extract of Buransh flower (AEBF)

The flowers of *Rhododendron arboreum* (Buransh) were collected from Mukteswar region of Nainital district of Uttarakhand. The plant material was authenticated at ICAR-Indian Veterinary Research Institute, Mukteswar, Uttarakhand, India and specimens were preserved in herbarium. The flowers were cleaned, air-dried, and pulverized in an electric grinder. The dried powder of Buransh flowers were subjected to aqueous extraction using water as solvent by columnar Soxhlet’s method at a temperature of 40-41°C with standard protocol from previous studies (Wang and Weller, 2006). Briefly, 15-20 gm of powdered material was taken in a thimble, 170-180 ml of distilled water was added in a 250 ml flask. Approximately, 6-8 cycles were repeated for complete extraction, as evidenced by the colourless liquid (aqueous extract of Buransh flower-AEBF) in the siphon tube of the thimble. The collected AEBF was dried using a vacuum evaporator for further analysis.

### *P. falciparum* growth inhibition assay

*P. falciparum* strains 3D7, Dd2 (chloroquine resistant) and C580Y (artemisinin resistant) cultures were revived from cryopreserved stocks and cultured in fresh A+ human erythrocytes with 10.4 g/L RPMI 1640 (Gibco), 5% sodium bicarbonate and Albumax II (Gibco) and supplemented with 50 mg/L gentamycin and 50 mg/L hypoxanthine; incubated at 5% CO_2_ and 37 °C. The parasite growth was monitored by Giemsa staining of thin smears. High parasitaemia cultures were synchronized using 5% sorbitol and 0.5% synchronized ring stage parasites were added to 96-well plates with 2.5% haematocrit. The effect of different concentrations of AEBF (100 to 0.097 µg/ml) was assessed by parasite growth inhibition assay using SYBR Green I fluorescent dye (Upadhyay et al., 2024). The experiment was performed in duplicate with three different biological replicates.

### Cytotoxic evaluation of AEBF

HepG2 Human Hepatocellular Carcinoma (HCC) cells were cultured in Dulbecco’s minimal essential medium (DMEM; Gibco, Thermo Fischer Scientific, USA) supplemented with 10% heat-inactivated fetal bovine serum (FBS; Gibco, Thermo Fischer Scientific, USA), and penicillin (10,000 Units/mL) and streptomycin (10,000 μg/mL) (Gibco, Thermo Fischer Scientific, USA). The cell line was maintained in a 25 cm^2^ culture flask at 37 °C in a humidified atmosphere of 5% CO_2_. Percentage cell viability was assessed using MTT (3-(4,5-Dimethylthiazol-2-yl)-2,5-Diphenyltetrazolium Bromide) assay (Ciapetti et al., 1993). Briefly, the cells were seeded in 96 well flat-bottom plates (1 x 10^4^ cells/well) for 24 hrs. The cells were then treated with AEBF at a concentration range from 7.81 µg/ml to 1000 µg/ml for 48 hrs. Post incubation, 10 μl of the MTT labelling reagent (final concentration 0.5 mg/ml) was added to each well containing 100 μl of cell suspension. The cells were further incubated for 4 hrs. After incubation, the media from each well was removed and 100 μl/well of DMSO was added and mixed gently. The absorbance was measured at 570 nm using a multi-well microplate reader (Varioskan Lux, Thermo Fischer Scientific, USA). The percentage viability of cells was calculated using the equation-% cell viability = [(Absorbance Treated)/ (Absorbance Control)] x 100. The experiment was performed with two different biological replicates.

### Therapeutic efficacy of AEBF against *P. berghei* infected mice

Adult male/female BALB/c mice (20 ± 3g of body weight) were used for the antimalarial *in vivo* testing and fed with water and food *ad libitum*. Experiments were performed according to the legal and ethical provisions including CPCSEA/IAEC guidelines and animal welfare regulations. The study was approved by the Institutional Animal Ethics Committee (IAEC) of ICMR-NIMR, New Delhi (IAEC/NIMR/2023-1/06). To assess the therapeutic efficacy of AEBF against *P. berghei* infected mice, 4 day Peter’s suppressive assay was followed (Peters, 1965; Upadhyay et al., 2024). Briefly, 0.2 ml of 1 × 10^5^ *P. berghei* NK65 pRBCs were injected into BALB/c mice intra-peritoneally. After visible asexual parasitemia on day 3 post inoculation, infected mice were divided into four groups and AEBF was administered as follows-IV: 50, 100 and 150 mg/kg); Oral: 200 mg/kg and two untreated mice were used as control. For survival analysis, AEBF treated groups and untreated controls were monitored until mice succumbed to infection. Mice were administered with intravenous AEBF doses twice per day, oral AEBF doses were administered once daily; both doses were administered for 4 consecutive days. Parasitemia was observed on alternate days by assessing Giemsa-stained thin smears made from a drop of blood from tails of mice (Rocha e Silva et al., 2011; Rocha e Silva et al., 2015).

### Assessment of AEBF in combination with Artesunate (ART)

*P. berghei* NK65 infected mice were divided in four groups (n=3 each)-1) Artesunate (ART) (dosing as per manufacturer’s protocol-FALCIGO, Zydus Cadila, India); 2) AEBF (Oral: 200 mg/kg for 4 days); 3) ART + BE; 4) Untreated controls and two mice were taken as non-infected controls. Protocol for administration of Artesunate was as follows: IV dose of 3mg/kg of body weight; 1st dose at 0 hrs, 2^nd^ dose at 12 hrs and final dose at 24 hrs. Parasitaemia was assessed by taking 2 µl blood from tail and Giemsa-stained thin smears were observed using oil immersed 100X objective in a compound microscope. Untreated mice were infected with the parasite *P. berghei* but did not receive any treatment whereas uninfected group were not given any infection. The doses were given according to the Peter’s test (Rocha e Silva et al., 2011; Rocha e Silva et al., 2015). Infection in the treated groups of Artesunate (3 mg/kg) and AEBF (Oral: 200 mg/kg) in combination was observed to have 100% inhibition of the parasite, and thus were re-infected with the parasites to check for survival in the two groups of mice (ART, ART + AEBF combination) without additional treatment.

### Investigation of splenomegaly and serum cytokines in *P. berghei* infected mice

Briefly, 0.2 ml of 1 × 10^5^ *P. berghei* NK65 pRBCs were injected into BALB/c mice intra-peritoneally. After visible asexual parasitemia on day 3 post inoculation; treated group of mice (n=3) were administered doses of AEBF+ART; while three control mice (n=3) received no treatment. One non-infected mouse was used as control. All the mice were sacrificed on the 7^th^ DPI; spleens were collected for further analysis. The mice were sacrificed using the cervical dislocation method under anaesthesia. Serums from the different groups of mice (AEBF treated, untreated and uninfected) were assessed for cytokine measurement. After 7 DPI, blood was collected from the mice’s heart and left for coagulation for 30 mins at 37 ^°^C. The samples were then centrifuged to obtain serum and stored at -80 ^°^C for further use. Cytokine levels in serum of mice were then determined using ELISA MAX (Biolegend™) cytokine kits for IFN-γ, TNF-α and IL-10 according to manufacturers’ protocol.

### Assessment of the transmission blocking potential of AEBF in *P. berghei ex vivo*

One group of mice (n=2) was treated with oral AEBF dose (200 mg/kg) for four days according to Peter’s test and another group (n=2) was taken as untreated controls. On the 10 days post infection (DPI), mature gametocytes were observed in *P. berghei* NK65 infected mice and percentage gametocytaemia was determined by Giemsa-stained thin blood smears. With similar gametocytaemia (∼1-1.2%) in both treated and untreated mice, blood was collected from the mice’s tails. To assess the reduction in the exflagellating centres, mature stage gametocyte containing blood (20 µl) from both groups of mice was mixed with 30 µl of ookinete medium (RPMI medium with 25mM HEPES, 20% human serum, 100μM xanthurenic acid, 50 mg/L hypoxanthine, 2 g/L sodium bicarbonate). Mixed blood was incubated at 19 °C for 15 minutes. The samples were then placed on a glass slide with a cover slip and observed under 100X objective in oil immersion. The fields in the microscope were then observed for exflagellation centres for the next 10-12 minutes and the no. of exflagellation centres were counted for both groups. The experiment was performed with two biological replicates.

### GC-MS analysis

GC-MS analysis of the AEBF was carried out at the Advanced Instrumentation Research Facility, JNU, New Delhi, using Shimadzu GC MS-QP-2010 plus system equipped with fused silica, RTx-5 SilMS column (30 m in length, 250 µm in diameter, 0.25 µm in thickness). Prior to analysis, a two-stage chemical derivatization process was performed in 100 µL of Methoxy amine hydrochloride (prepared in pyridine at 20 mg/mL concentration) followed by vortex mixing for 2-5 min and subsequent incubation at RT for 1 hr to convert aldehyde and ketones into their oximes or alkyloximes. Then, 100 µL of N-methyl-N-trimethylsilyltrifluoro(o) acetamide was added to this mixture in 1:1 ratio and incubated at 70 °C for 30 min to make the metabolites in the mixture less polar and more volatile, before the analysis. Different components present in the test samples were identified based on the comparison of their retention time (min), peak area, peak height, and mass spectral patterns with those of the spectral database of authentic compounds stored in the National Institute of Standards and Technology (NIST) and WILEY library.

### Molecular docking analyses of AEBF bioactive constituents and putative protein targets in *P. falciparum*

#### Ligand preparation

The 2D structures of the candidate phytochemicals, erythritol (CID 222285), oleic acid (CID 445639), quercetin (CID 5280343) and rutin (CID 5280805) were retrieved from the PubChem database (Kim et al., 2022) and prepared in the LigPrep module of the Schrödinger Suite (Schrödinger Release 2023-2; Schrödinger, LLC, 2023) (Jacobson et al., 2017) using the OPLS4 force field (Lu et al., 2021). At pH 7.0 ± 2.0, the number of ionisation and tautomeric states were enumerated using Epik (Johnston et al., 2023), up to 32 stereoisomers per ligand were generated, containing input chiralities.

#### Target preparation

Five potential targets for *P. falciparum* were chosen from literature sources **(Table 2)**. The crystal structures of aquaglyceroporin (PDB 3C02), plasmepsin I (PDB 3QRV) and apical membrane antigen 1 (AMA1; PDB 3SRI) were downloaded from the PDB (Berman et al., 2000; Burley et al., 2023). The models of ferredoxin (UniProt Q8ILM8) and the small heat shock protein PfHsp20a (UniProt Q8IES0; with an additional entry for the small heat shock protein of *P. falciparum*, A0A0L1I6A3) were downloaded from the AlphaFold Protein Structure Database through UniProt (Jumper et al., 2021; The UniProt, 2023; Varadi et al., 2024). The structures were prepared in the Protein Preparation Wizard (Sastry et al.) and processed as follows, prior to processing each target: Bond orders were assigned to the structures, hydrogens were added, and the ionisation states were determined using the pH 7.0 addition method of PROPKA (Olsson et al.). The structure was then subjected to an OPLS4 restrained minimisation until the heavy-atom RMSD reached 0.30 Å. Waters farther than 5 Å from het groups were removed.

#### Binding-site prediction and grid generation

The putative ligand binding pocket was predicted by using SiteMap (Halgren), and only sites with SiteScore ≥ 0.80 were kept. Some targets returned one qualifying pocket, while others returned two - the pockets that met the cut-off were each carried forward, one pocket to be treated as a single receptor grid, with the Receptor Grid Generation panel of Glide, using the OPLS4 force field and no constraints (default 10 × 10 × 10 Å inner box).

#### Molecular docking using Glide-XP

Flexible docking was carried out using the Extra Precision (XP) mode of Glide (Friesner et al.), with default settings (0.80 for van der Waals scaling, 0.15 e for partial-charge cut-off, Epik state penalties were applied, and post-docking minimisation was turned on). A total of one Glide-XP run per pocket per ligand/fold (e.g., one run per pocket for one ligand vs. two runs (P1 and P2) for two ligands/folds) was performed for each ligand/target pair. The complexes have been ranked on the basis of the Glide docking score and Glide-XP GScore with negative values representing more negatively predicted binding (Friesner et al.). The Ligand Interaction Diagram tool of Maestro was used to visualise the protein–ligand interactions.

## Results

### In vitro antimalarial evaluation of AEBF against human parasite-*P. falciparum*

To assess the in vitro antimalarial potential of AEBF, *P. falciparum* asexual stage parasites were exposed to various concentrations of AEBF, in vitro. Microscopic examination of the thin blood smears revealed significant inhibition of the parasite growth at both 50 and 100 µg/ml concentrations, with complete parasite clearance at higher concentration, comparable to the positive control chloroquine (CQ) **(Fig. 1A)**. Further, serial dilutions of AEBF (100 to 0.097 µg/ml) were tested against *P. falciparum* 3D7 and Dd2 (chloroquine resistant) strains, yielding an IC_50_ of 16.2 µg/ml and 29.4 µg/ml, respectively **(Fig. 1B & 1C)**. The growth inhibitory potential of AEBF against artemisinin resistant strain *P. falciparum* C580Y revealed an IC_50_ value of 15.2 µg/ml **(Fig. 1D)**. The resistance index (RI) of both the drug-resistant strains was observed to be in the range of 1-2, indicating their low resistance potential against AEBF **(Fig. 1C & 1D)**.

**Figure 1:**
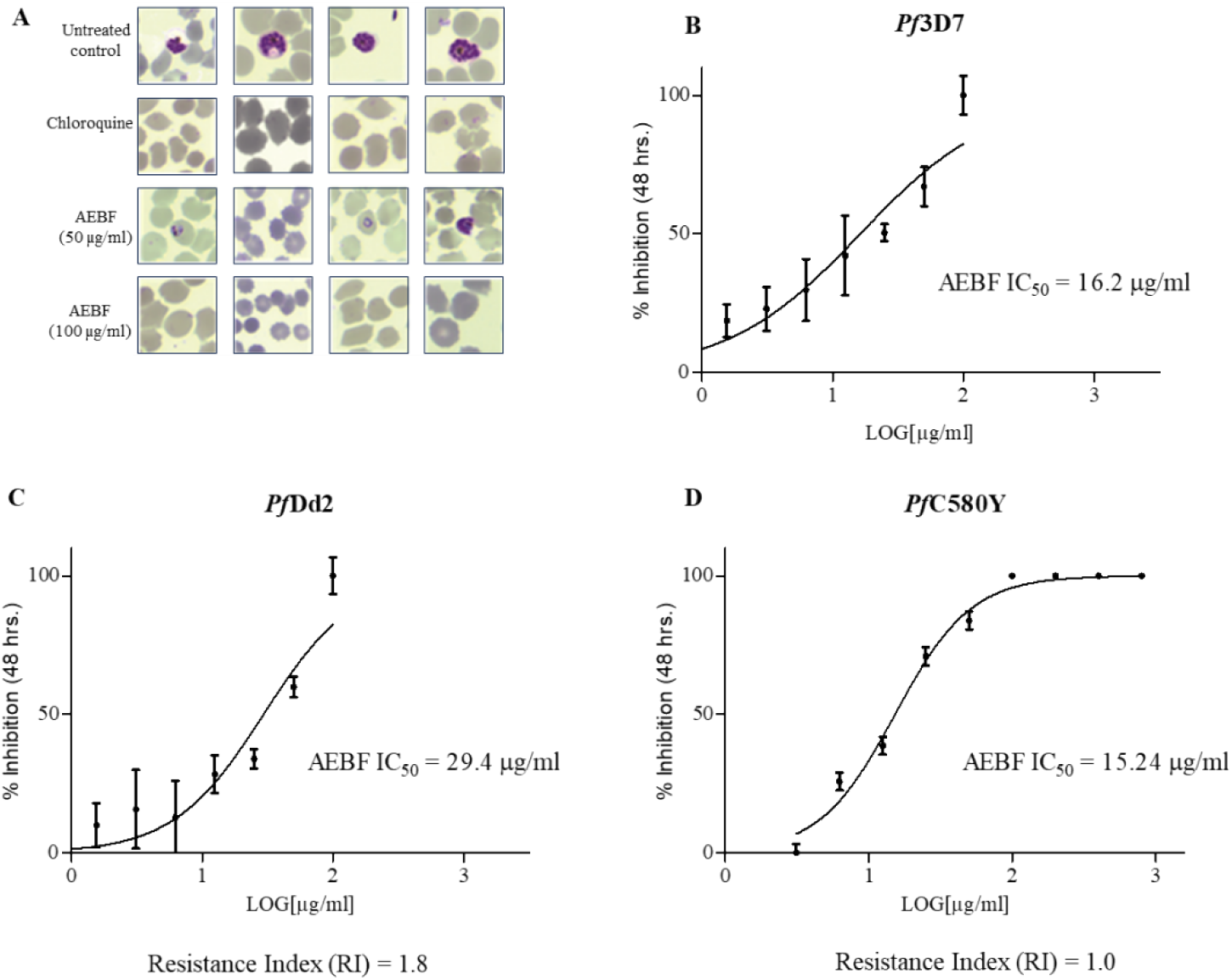
Therapeutic potential of Aqueous Extract of Buransh Flower (AEBF) against asexual stages of various *P. falciparum* strains. **A)** Giemsa-stained images of *P. falciparum* cultures after 48 hrs of AEBF exposure-Untreated control, Chloroquine (CQ)-positive control, AEBF (50 µg/ml) and (100 µg/ml). **B)** IC_50_ estimation of AEBF against *P. falciparum* 3D7 strain indicated by % inhibition of the parasite growth **C)** IC_50_ estimation of AEBF against *P. falciparum* Dd2 (chloroquine resistant) strain indicated by % inhibition of the parasite growth. **D)** IC_50_ estimation of AEBF against *P. falciparum* C580Y (artemisinin resistant) strain indicated by % inhibition of the parasite growth. Resistance Index (RI) of AEBF against *Pf*Dd2 and *Pf*580Y strains is indicated in panels C and D.

### Cytotoxicity assessment of AEBF in human cell line-HepG2 HCC

To assess any detrimental impact of AEBF on human cells, we investigated its cytotoxicity on HepG2 hepatocellular carcinoma (HCC) cell line using the MTT (3-(4, 5-dimethylthiazolyl-2)-2, 5-diphenyltetrazolium bromide) cytotoxicity assay. The extract was tested across a concentration range from 7.81 µg/ml to 1000 µg/ml. No significant effect on cell viability was observed up to 1000 µg/ml, indicating that AEBF does not exhibit any toxicity towards human cell line HepG2 HCC **(Fig. 2)**. Interestingly, AEBF appeared to promote cell proliferation, in a dose-dependent manner **(Fig 2)**.

**Figure 2:**
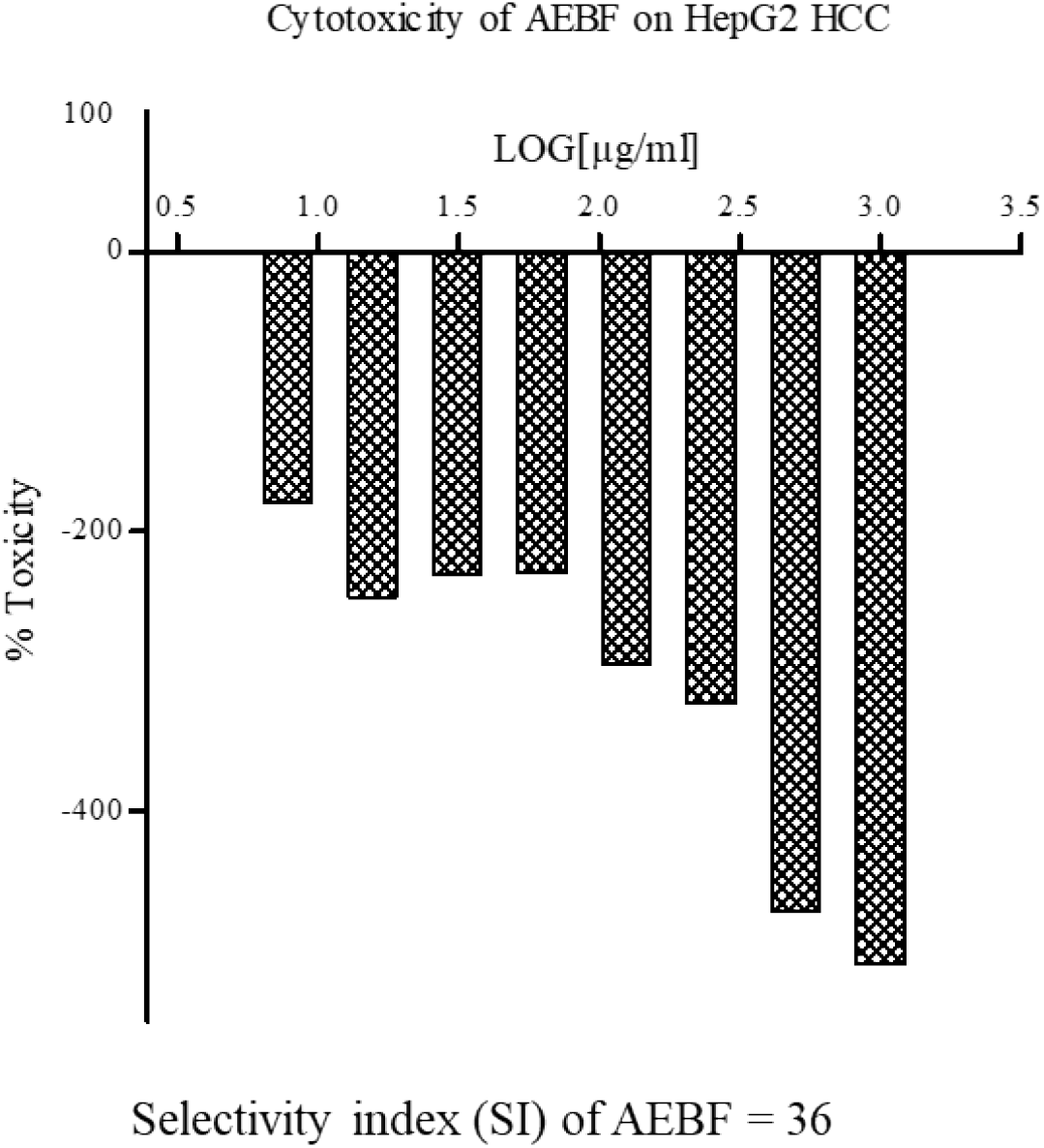
Cytotoxicity evaluation of AEBF on HepG2 HCC cell line. Bar graph showing % Toxicity using MTT assay in HepG2 HCC cell line. All the tested doses (7.81 µg/ml to 1000 µg/ml) demonstrated no toxicity; instead, the proliferative effect at higher doses was clearly evident. The % Toxicity was tested in triplicates for all tested concentrations.

### In vivo antimalarial evaluation of AEBF in *P. berghei* infected mice

To evaluate the antimalarial potential of AEBF, we performed a 4-day Peter’s suppressive assay in BALB/c mice infected with *P. berghei* NK65 (Peters, 1965). Mice were divided into four groups and administered varying doses of AEBF (IV: 50, 100, and 150 mg/kg and Oral: 200 mg/kg of body weight) with two untreated mice serving as controls **(Fig. 3A)**. All IV doses of AEBF reduced parasitemia (63-80 %) on 7^th^ day post-infection (DPI) and extended the mean survival time (MST) from 13 days for untreated control to 18-19.5 days for IV treated mice **(Fig. 3B & 3C)**. Interestingly, the oral dose of AEBF resulted in 51% reduction in asexual parasitemia compared to untreated controls **(Fig 3B)**; but extended the survival of mice up to 23.5 days **(Fig. 3C)**. Representative Giemsa-stained thin smears of AEBF oral treated and untreated controls are shown in **Fig. 3D**. To further investigate the marked extended MST in spleen size, we compared the spleen size of orally administered AEBF and untreated controls. A significant reduction in spleen size was observed in AEBF treated mice, potentially by modulation of splenic parasite clearance, maintaining the natural architecture of spleen **(Fig. 3E)**. Given the maximum enhancement in the MST by oral dose of AEBF and its palatability owing to its natural sugar content, subsequent experiments focused on the oral route of administration.

**Figure 3:**
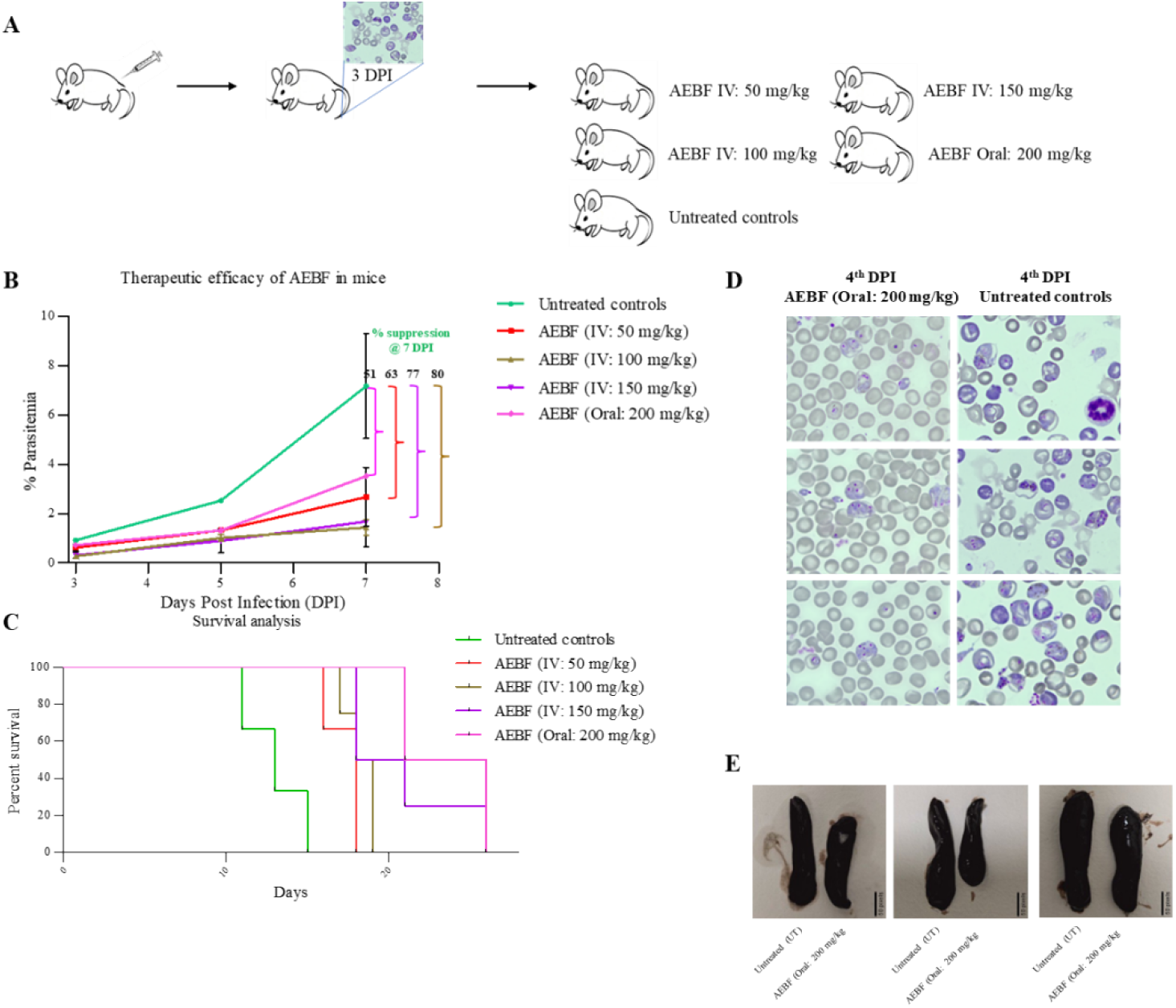
Assessment of the antimalarial effects of AEBF on *P. berghei* infected mice. **A)** A schematic showing the various groups of BALB/c mice infected with *P. berghei* NK65 and the doses of AEBF received and untreated controls. **B)** % parasitemia in the infected mice AEBF [IV: 50-red; 100 brown & 150-violet and Oral: 200-pink); untreated mice-green. **C)** % survival of AEBF treated mice (IV: 50-red; 100-brown and 150-violet; Oral: 200-pink and untreated controls-green). **D)** Representative Giemsa-stained blood smears from tail blood of AEBF Oral (200 mg/kg) treated mice and untreated control mice. **E)** Pictures of spleen sizes of untreated controls and AEBF Oral (200 mg/kg) treated mice to show enlargement of spleen on 7^th^ DPI with scale bar.

### Evaluation of AEBF combined with Artesunate and immunomodulation in *P. berghei* infected mice

To evaluate the synergistic effects of AEBF with frontline antimalarial therapy, we tested its combination with Artesunate (ART) in *P. berghei*-infected mice. Mice were divided into four groups (n=3 each): 1) ART (3 mg/kg, intravenous, 3 days); 2) AEBF (200 mg/kg, oral, 4 days); 3) ART+AEBF; and 4) untreated controls; two non-infected mice served as additional controls. Both ART and ART+AEBF combination treatments achieved 100% parasite clearance, while AEBF alone resulted in 55% reduction in asexual parasitemia on 8^th^ DPI compared to controls **(Fig. 4A)**. Survival analysis showed extension of MST up to 27 days for ART and ART+AEBF and 20 days for AEBF alone **(Fig. 4B)**. Parasite reduction in all the treated mice was evident by Giemsa-stained thin smears **(Fig. 4C)**. Importantly, there was a visible reduction in splenomegaly for ART+AEBF treated mice, compared to ART alone **(Fig. 4D)**. Further, to assess sustained protection, mice previously cleared of parasites by ART alone and ART+AEBF were reinfected, without further treatment. Notably, the ART+AEBF combination exhibited a marked reduction (67%) in parasitemia, on day 7^th^ DPI compared to ART alone **(Fig. 4E)**. To investigate the potential immunomodulation, we assessed serum levels of three cytokines: IFN-γ, TNF-α, and IL-10 on 7^th^ DPI, in both AEBF treated and untreated mice. A significant elevation was observed in serum IFN-γ levels in AEBF treated mice compared to untreated controls (p-value = 0.0204) **(Fig. 4F)**. While both TNF-α and IL-10 levels were elevated in AEBF treated mice, the differences were not statistically significant **(Fig. 4F)**. Altogether, these findings strongly indicated immunomodulation by AEBF resulting in sustained protection even in cases of reinfection, without the need for further treatment.

**Figure 4:**
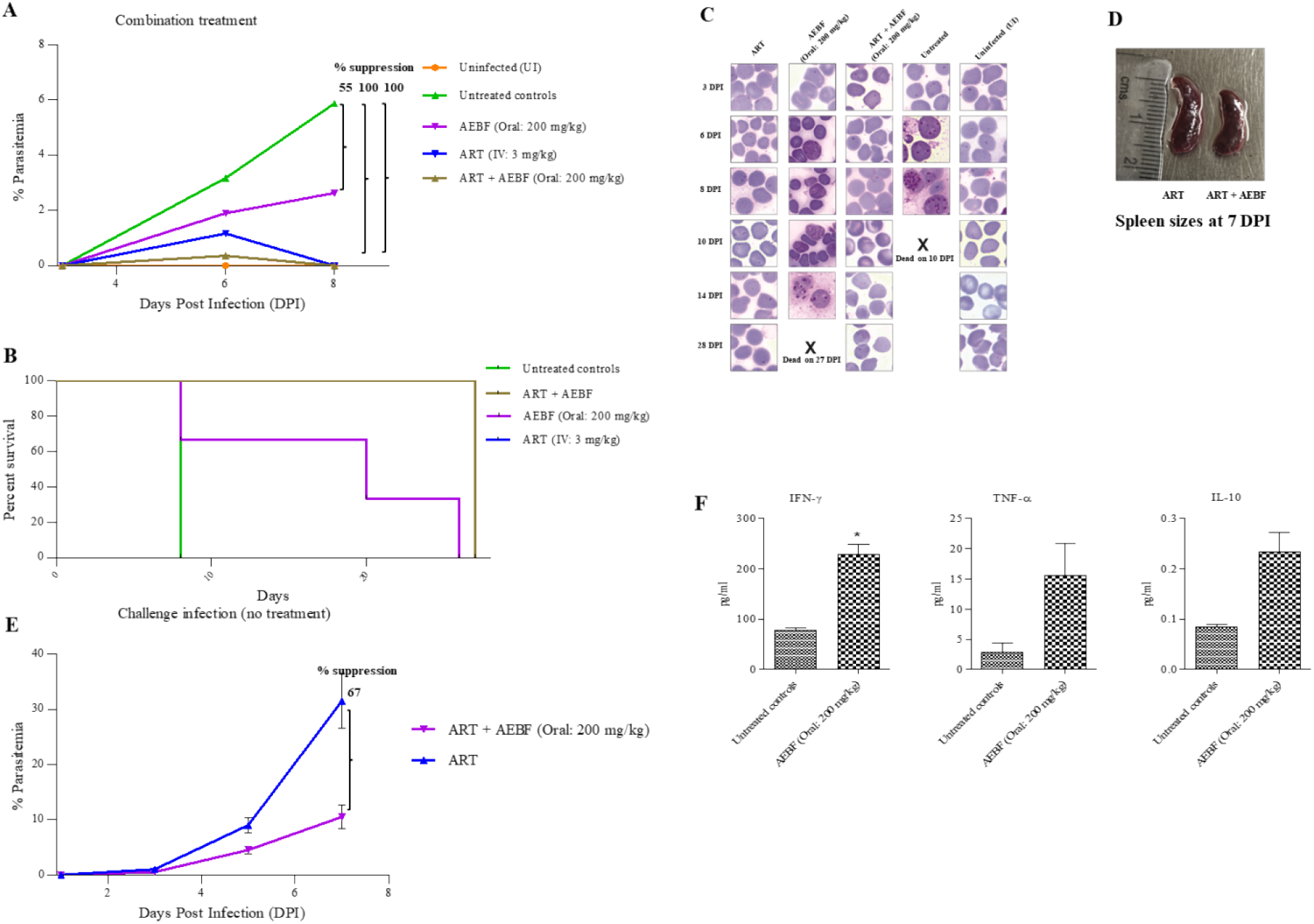
Combination treatment of AEBF with artesunate (ART) in *P. berghei* infected mice and potential immunomodulation. **A)** % parasitemia of five experimental groups in primary infection (Uninfected (orange); Untreated (teal); AEBF (violet); ART + AEBF (green) and ART (blue). **B)** Survival analysis of the ART, BE, ART + BE and untreated control groups. **C)** Giemsa-stained thin blood smears of from five groups of mice-(ART, AEBF, ART + AEBF, Untreated controls, Uninfected (UI)). **[X]** represents the mice died before checking asexual parasitemia on respective DPI. **D)** Pictures of spleens from ART and ART + AEBF treated mice upon challenge at 7^th^ DPI. **E)** Bar graphs showing the levels of serum cytokines-IFN-γ, TNF-α and IL-10 in mice on 7^th^ DPI. The significant value is represented by (*). **F)** % parasitemia in challenge infection in treated mice from groups-ART and ART + AEBF; without new treatment. Significance was estimated using two-tailed t-test in Graphpad Prism.

### AEBF treatment reduced exflagellation of male gametocytes in *P. berghei*

Counting exflagellating centres is a reliable indicator of the functional viability of mature male gametocytes-sexual stages in mosquito vector highlights the transmission-blocking potential of antimalarials (Ruecker et al., 2014). Blood from AEBF treated mice (oral: 200 mg/kg) containing sexual stages-mature gametocytes was incubated with ookinete media and the number of exflagellating centres were compared to that from controls **(Fig. 5A)**. AEBF treated samples showed a marked reduction in the exflagellating centres; 25±9 in blood of AEBF treated mice versus 71±18 in control mice **(Fig. 5B)**. It is important to note that the gametocytaemia levels were similar between both groups (data not shown), indicating that the reduction was specific to the male gametocyte exflagellation rather than a general effect on parasite sexual development.

**Figure 5:**
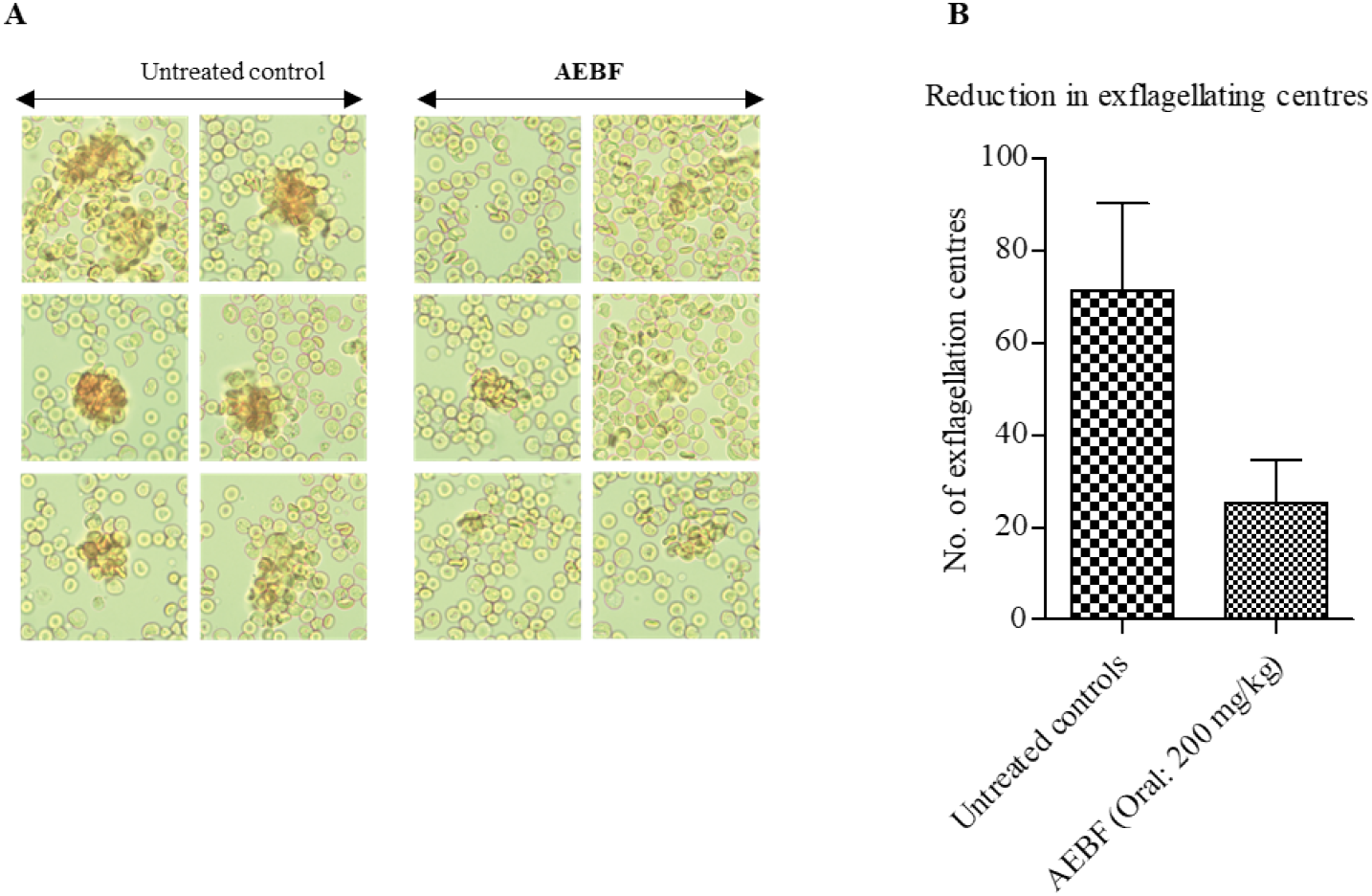
Reduction in exflagellation centres from male gametocytes of *P. berghei*. **A)** Left panels showing exflagellating male gametocytes forming centres on glass slide. Right panel showing reduced exflagellating centres from BE treated mice. **B)** Bar graphs showing reduction in exflagellating centres counted on glass slide from untreated controls and BE treated mice. Two separate experiments were performed with two different biological replicates.

### Evaluation of the pleiotropic targets of AEBF bioactives in *P. falciparum* via molecular docking

Gas chromatography-mass spectrometry (GC-MS) analysis of AEBF revealed a complex phytochemical profile comprising of 86 distinct compounds including sugars, sugar alcohols, organic acids and other secondary metabolites **(Fig. S1** and **Table S1)**. Quantitative profiling deduced from the relative composition in AEBF indicated that quininic acid was the most abundant constituent, followed by various sugars (D-fructose, D-(+)-Talose and sucrose) and fatty acids including palmitic acid and oelsaeure (Oleic acid) **(Fig. 6)**. To elucidate the potential antimalarial mechanisms of these constituents, selected bioactive compounds were docked against *P. falciparum* protein targets, carefully selected from literature review **(Table 1)**. All Glide-XP docking jobs were completed successfully with binding energies ranging from −7.86 kcal mol^−1^ (quercetin-PfHsp20a (Q8IES0)) to −4.41 kcal mol^−1^ (quercetin-PfHsp20a (A0A0L1I6A3), pose P2). The thermodynamic stability and interaction profiles of the top ranked poses are summarized in **Table 2**. Quercetin demonstrated high binding affinity to PfHSP20a (Timothy and Zininga, 2025). It achieved the highest overall docking score of −7.86 kcal mol^−1^ against the AlphaFold model of PfHsp20a (Q8IES0) **(Fig. 7)**. The score of the alternate PfHsp20a entry A0A0L1I6A3 was −6.996 kcal mol^−1^ (GScore −7.036 kcal mol^−1^) for pocket P1 and −4.416 kcal mol^−1^ (GScore −4.455 kcal mol^−1^) for pocket P2, showing clear pocket selectivity. Rutin displayed strong binding to the apical membrane antigen 1 (AMA1). Pocket P2 yielded the strongest docking score of −7.83 kcal mol^−1^. Notably, pocket P3 exhibited the study’s most significant H-bond contribution (−4.37 kcal mol^−1^) and Coulombic interaction (−20.41 kcal mol^−1^), driven by extensive polar contacts between the rutinose moiety and the protein surface **(Fig. 8)** (Bai et al., 2005). Erythritol exhibited a docking score of −4.59 kcal mol^−1^. The interaction was primarily driven by hydrogen bonding between the ligand’s polar tetraol scaffold and the channel’s substrate-binding site **(Fig. S3)** (Beitz et al., 2004). The relative affinities of the ligands were ranked based on their most stable docking poses **(Table 2)**: quercetin–PfHsp20a (−7.86 kcal mol^−1^) > rutin–AMA1 (−7.83 kcal mol^−1^) > quercetin– PfHsp20a (A0A0L1I6A3, −7.00 kcal mol^−1^) > erythritol–aquaglyceroporin (−4.59 kcal mol^−1^). Given that quercetin and rutin consistently produced docking scores below the −7 kcal mol^−1^ threshold-a value generally indicative of strong predicted binding in Glide-XP. These flavonoids could be explored as promising lead compounds for further experimental validation of their antimalarial potential.

**Table 1:** List of important bioactives in AEBF and their putative protein targets in *P. falciparum*.

| S. no. | Compound | Putative protein target |
| --- | --- | --- |
| 1. | Quercetin (CID 5280343) | Small Heat Shock Proteins-<br>*PfHsp20a*- UniProt Q8IES0 (Timothy and Zininga, 2025) |
| 2. | Rutin (CID 5280805) | AMA1- PDB 3SRI (Dangana et al., 2026) |
| 3. | Erythritol (CID 222285) | Aquaglyceroporin- PDB 3C02 (Kumari et al.) |

**Table 2:**
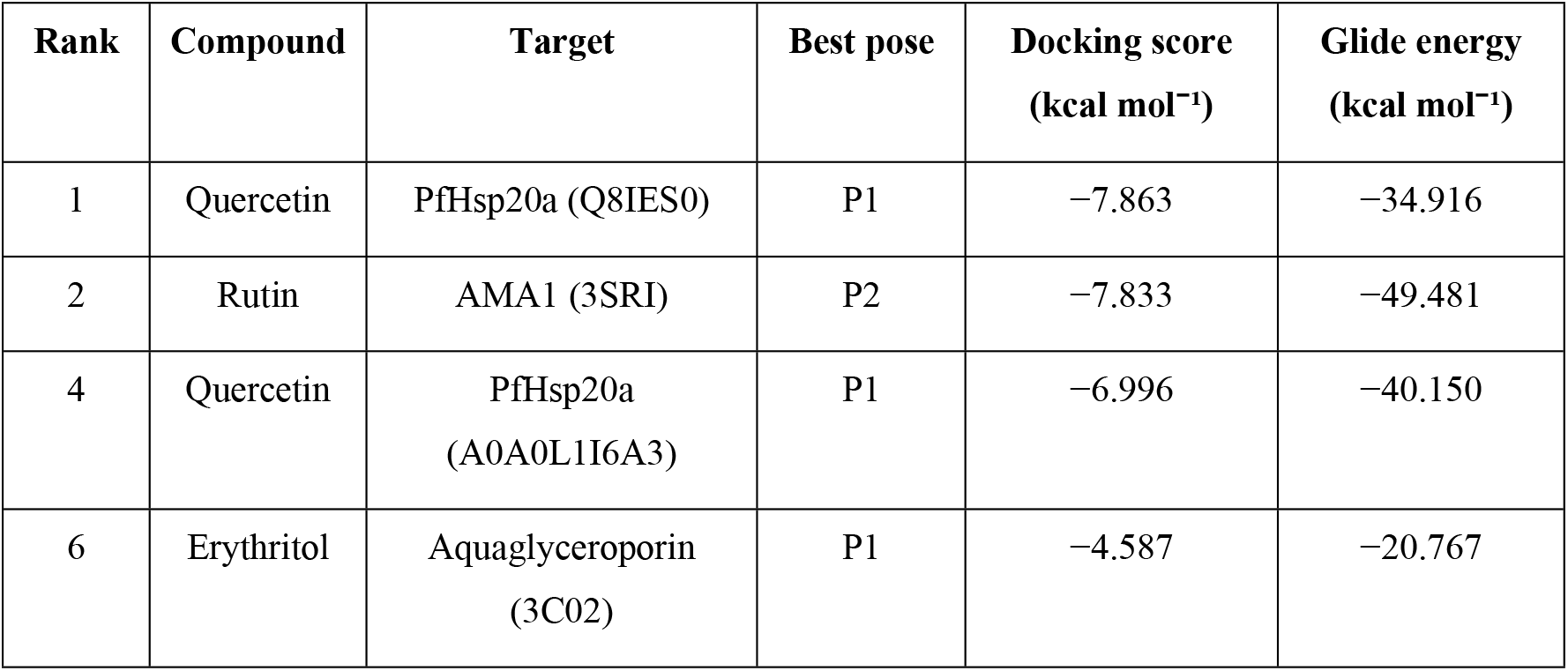
Best-scoring Glide-XP pose retained for each ligand–target pair and its overall ranking within the screened set.

**Figure 6:**
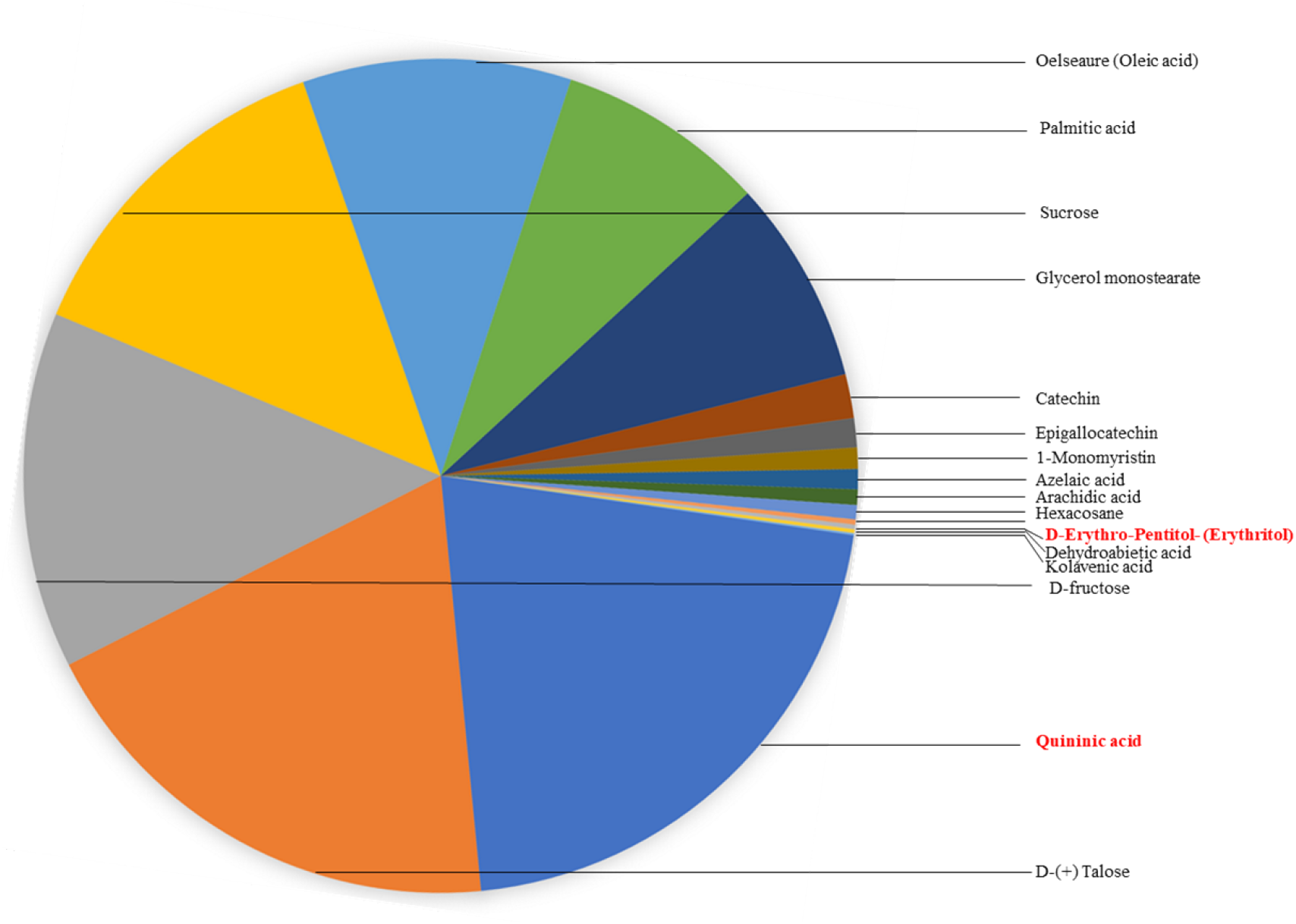
Relative composition of volatile mix of Aqueous extract of Buransh Flower (AEBF) in Gas Chromatography Mass Spectrometry (GC-MS) analysis.

**Figure 7:**
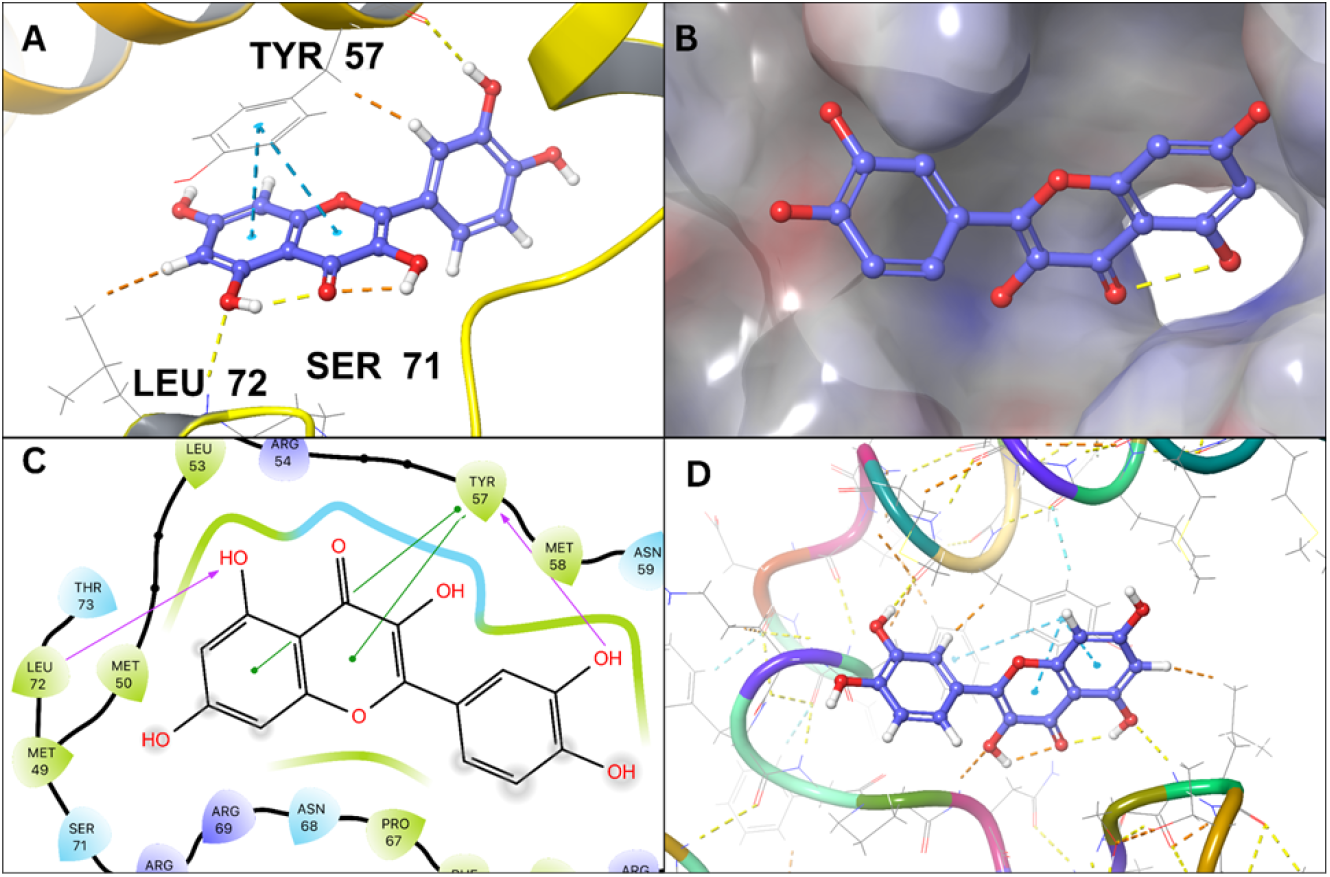
Glide-XP docked poses and ligand-residue interaction diagrams of quercetin in PfHsp20a (Q8IES0, AlphaFold)

**Figure 8:**
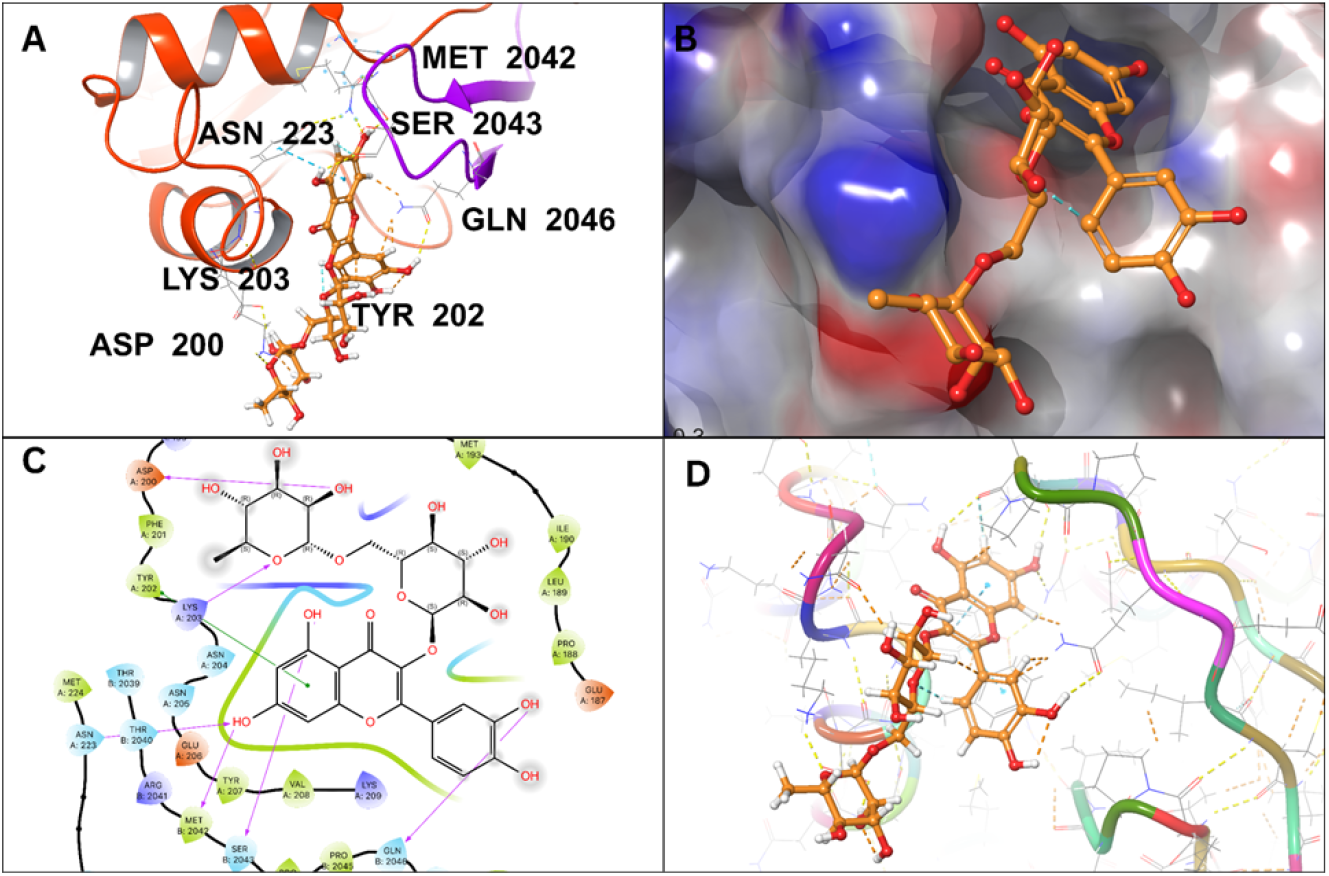
Glide-XP docked poses and ligand-residue interaction diagrams of rutin in pockets of AMA1 (PDB 3SRI).

## Discussion

This study evaluated the antimalarial potential of Aqueous extract of Buransh flower (AEBF) in human and rodent malaria using in vitro cell studies, in vivo therapeutic evaluation and molecular docking of buransh flower bioactive constituents against selected *P. falciparum* proteins. The observed pleiotropic antimalarial activity of AEBF likely stems from its complex nature of bioactive molecules, obtained from GC-MS analyses and review of existing literature on bioactives of *R. arboreum* flowers. Quininic acid, a metabolic by-product of the frontline antimalarial quinine, represents the most abundant constituent in AEBF; while it lacks direct antimalarial activity, it serves as a critical scaffold for the synthesis of quinine analogues and other 4-quinolinecarboxylic acid derivatives (Verganista et al., 2026). The literature review of bioactives in Buransh flower reported the presence of two phenolic compounds-quercetin and rutin by HPTLC analysis (Swaroop et al., 2005). Quercetin has been shown to disrupt the structural integrity of the small heat shock proteins (sHSPs) of *P. falciparum* (PfHSP20a, PfHSP20b and PfHSP20c) in a dose-dependent manner. The in vitro antimalarial activity of quercetin was observed to be 5µM and 8 µM for *P. falciparum* NF54 (chloroquine sensitive) and Dd2 (chloroquine resistant) strains, respectively (Timothy and Zininga, 2025). Furthermore, Rutin has shown significant in vitro activity against both chloroquine-sensitive and resistant strains of *P. falciparum*, primarily through antioxidant properties, reducing oxidative stress and inflammation. Oral administration of rutin has been shown to reduce parasitaemia, increase mean survival time, and restore haemoglobin/glucose levels in *P. berghei*-infected mice (Bhatt et al.). Erythritol-a safe natural sweetener, previously demonstrated multifaceted antimalarial activity through disruption of parasitic hydro-homeostasis and cytokine modulation effect. Erythritol infiltrates the parasite potentially by acting as a permeant of the *P. falciparum* aquaglyceroporin (PfAQP) channel and induces significant osmotic stress, ultimately resulting in osmotic lysis. This disruption is effective across multiple life-cycle stages, inhibiting the development of sporozoites in the liver, asexual progression in the blood (including invasion and egress), and the maturation of gametocytes for transmission (Kumari et al., 2022). Interestingly, the IV administration resulted in improved parasite clearance than oral dose, while the latter conferred superior survival rates in the murine malaria model. We hypothesize that oral delivery facilitates a more sustained therapeutic window and potentially activates host-mediated immunomodulatory pathways-such as systemic cytokine regulation that alleviate malaria-associated pathology. Furthermore, the first-pass metabolism of AEBF constituents may yield bioactive metabolites with enhanced long-term protective effects, an observation that warrants further pharmacokinetic and pharmacodynamic investigation. Moreover, the MSTs achieved in mice with AEBF treatment were notably superior to those reported in previous studies using natural plant extracts (Adigo Shibeshi et al., 2021; Ha et al., 2015; Hagazy et al., 2020; Mekonnen, 2015).

Artemisinin-based Combination Therapies (ACTs) combine the rapid-acting drug artemisinin with longer-acting partner drugs such as sulfadoxine, pyremethamine or lumefantrine. Our study demonstrated a synergistic effect between AEBF and Artesunate (ART), achieving complete parasite clearance after primary infection. Notably, the AEBF+ART combination also led to a significant reduction in parasitemia upon *P. berghei* infection challenge, without additional treatment. These observations suggested the role of AEBF in inducing a sustained protective or memory immune response. The significant elevation of serum IFN-γ, a proinflammatory cytokine in orally treated mice would enhance the immune defense against intracellular parasites (Deng et al., 2023; Villegas-Mendez et al., 2012). IFN-γ mediates a delicate balancing act in malaria; while it is crucial for parasite clearance, excessive or prolonged IFN-γ drives immunopathology, including the breakdown of splenic architecture and severe splenomegaly. Inhibiting this excessive inflammation to preserve spleen function requires strategic modulation. Splenomegaly (enlargement of spleen) is a hallmark of malaria and plays a key role in clearing infected red blood cells (RBCs) by macrophages as well as facilitating the activation and proliferation of B & T cells, essential for targeting malaria parasites (Ghosh and Stumhofer, 2021; Nuss, 2005). Previous research by Ha et al. had shown that the isoflavone genistein can restore disrupted spleen architecture to a state similar to that of uninfected spleens (Ha et al., 2015). TNF-α, also contributes to protection against parasitic infections (Leão et al., 2020), while, IL-10, an anti-inflammatory cytokine, regulates the activity of both IFN-γ and TNF-α (Kumar et al., 2019; Tembo et al., 2023). Although a comprehensive cytokine profile would be needed to fully assess the immunomodulatory effects of AEBF against *P. berghei* infection, such analysis was beyond the scope of the present study. While these results are promising, more detailed histopathological analysis would provide greater insight into the specific structural and functional changes induced by AEBF. Further, clinical trials are necessary to evaluate the efficacy of this combination in patients with drug resistant malaria infections. We are currently collaborating with experts form the Greater Mekong region to address the challenges of artemisinin resistance and are planning prospective clinical trials. The ability of AEBF to affect the transmissible sexual stages of *P. berghei* is promising suggesting that its use as a health drink in endemic communities could help disrupt the malaria transmission cycle. This approach of using AEBF along with frontline antimalarials can be impactful for malaria elimination efforts in holoendemic regions such as sub-Saharan Africa.

## Conclusion

Altogether, our study has generated first ever evidence of the pleiotropic antimalarial and immunomodulatory properties of AEBF, using in vitro cell studies, in vivo therapeutic evaluation, further combined with molecular docking analyses of Buransh flower bioactive constituents with putative protein targets in *P. falciparum*. The absolute lack of cytotoxicity of AEBF on human cell lines, together with its traditional use as health drink in communities across Himalayan region of India, advocates for its safety for future clinical trials. Crucial findings of our study show that AEBF exhibits antimalarial activity against sensitive and resistant *P. falciparum* strains, prolonged survival, provided sustained immunity, prevented splenomegaly, and potentially disrupted transmission. These results highlight the potential of Buransh flower juice as an adjunct therapy for drug-resistant malaria, subject to thorough clinical evaluation.

## Abbreviations

AEBF: Aqueous extract of Buransh flower
*P. falciparum*: *Plasmodium falciparum*
*P. berghei*: *Plasmodium berghei*
HCC: HepG2 Human Hepatocellular Carcinoma
MTT: (3-(4,5-Dimethylthiazol-2-yl)-2,5-Diphenyltetrazolium Bromide)
IFN-γ: Interferon-γ
TNF-α: Tumor Necrosis Factor-α
IL-10: Interleukin-10
ART: Artesunate
RPMI-1640: Roswell Park Memorial Institute 1640
HEPES: 4-(2-hydroxyethyl) piperazine-1-ethanesulfonic acid

## Declarations

## Acknowledgements

We express our sincere gratitude to the following individuals and institutions for their invaluable contributions to this research:

## Technical Support

Dr. Nidhi Sharma for technical assistance in experiments.

Mrs. Bhavana Tilsari for collecting flowers from the Mukteswar region.

Members of the Animal House at ICMR-NIMR for their technical support.

## Ethical Clearance

Institutional Animal Ethics Committee (IAEC) and Committee for the Purpose of Control and Supervision of Experiments on Animals (CPCSEA, Registration No. 33/GO/ReBi/S/99/CSCSEA) for approving ethical clearance for the use of mice in in-vivo experiments (IAEC/NIMR2023-1/06).

The publication committee of ICMR-National Institute of Malaria Research, New Delhi, for approving the manuscript.

## Clinical trial number: not applicable

Council of Scientific and Industrial Research (CSIR) for providing fellowship and Academy of Scientific and Innovative Research (AcSIR) for the doctoral program (10BB20A65074) for Ms. Cherish Prashar.

## Author contributions

Conception: KCP, AG, KV

Investigation: CP, NT, RST, SA, RH, RB, AM, PR, HLS, PRB, HK

Manuscript writing: CP, KV, SA, KCP

Manuscript review: KCP, KV, AG, SS, MA, JD, BR, RKJ, SC, VS

## Author statement

All authors have seen and agree with the contents of this manuscript and the journal policies.

## Declaration for use of generative AI technology

During the preparation of this work the author(s) used Gemini AI platform in order to achieve minor language enhancement. After using this tool/service, the author(s) reviewed and edited the content as needed and take(s) full responsibility for the content of the publication.

